# A framework for detecting and quantifying relationship alterations in microbial community: Quantifying microbial relationship alteration

**DOI:** 10.1101/2020.04.09.033688

**Authors:** Zhi Liu, Kai Mi, Zhenjiang Zech Xu, Qiangkun Zhang, Xingyin Liu

## Abstract

Dysbiosis of gut microbiota is associated with the pathogenesis of human disease. Observing shifts in the microbe abundance cannot fully reveal underlying perturbations. Examining the relationship alteration (RA) in microbiome between different healthy status provides additional hints about the pathogenesis of human disease. However, no methods were designed to directly detect and quantify the RA between different conditions. Here, we present PM2RA (Profile Monitoring for Microbial Relationship Alteration), an analysis framework to identify and quantify the microbial RAs. The performance of PM2RA were evaluated in synthetic data, and found to show higher specificity and sensitivity than the co-occurrence-based methods. Analyses of real microbial dataset show that PM2RA is robust for quantifying microbial RA across different datasets in several diseases. By applying PM2RA, we identified both previously reported and novel microbes implicated in multiple diseases. The PM2RA is implemented as a web-based application available at http://www.pm2ra-xingyinliulab.cn/.

## Introduction

The gut microbiome is considered to be the second genome and is linked to many human diseases[1–3]. A central goal of human microbiome studies is to identify microbes associated with disease. The identified microbes can provide insights into disease etiology as well as having the potential for use as biomarkers for disease diagnosis and prevention, and if causal, as therapeutic targets.

The development of next-generation sequencing technologies enables culture-independent investigations of the role of the human microbiome in health and disease via direct DNA sequencing. Both 16S rRNA sequencing and metagenomic sequencing have been used to study the human microbiome, allowing the creation of an abundance table based on which differential analysis between different biological conditions can be performed[4–6]. Although differential abundances of certain microbes may contribute towards conferring a specific trait in a given condition, alterations in the level of a single microbe have limited capacity to reflect the real perturbation of ecological networks under phenotype change.

The human microbiome is a complex bacterial community that forms sub-communities based on shared niche specializations and specific interactions between individual microbes. The mutual associations within the residing microbial communities play an important role in the maintenance of eubiosis [7–9]. Bacteria can interact with each other in numerous ways, such as commensalism, mutualism, and competition, which can cause beneficial, neutral, or detrimental effects for the microbes involved. Commensalism refers to situations where some constituent microbes of an ecosystem derive benefit from other members without helping or harming them, while mutualism describes to interactions between microbes where both organisms benefit from each other[8]. A microbe might also directly compete with another for the same nutrition source, thereby creating a competition[7,10]. These kinds of functional relationships are referred to *profiles* which can be linear, polynomial, nonlinear or a waveform[11]. The disruption of these relationships can lead to disorders in the microbial community structure, furthering dysbiosis.

The relationships are usually modeled by the linear correlation between two types of microbes [12–14], and co-occurrence networks are constructed to describe the whole microbial relationship. Based on the co-occurrence networks, alignment-based[15–19] or alignment-free[20–22] methods have been proposed to visualize the relationship changes between different conditions, e.g. health vs. disease. The alignment-free network comparison methods aim to quantify the overall topological similarity between networks, irrespective of node mappings between the networks and without aiming to identify any conserved edges or subgraphs, and the alignment-based methods aim to find a mapping between the nodes of two networks that preserves many edges and a large subgraph between the networks. These strategies are unable to quantify the association change for a specific group of microbes or pinpoint the exact nodes that contribute to the community difference between two conditions.

Applying the concept of profile monitoring which is widely used to monitor the relationship consistency between variables in the industry of food production, manufacturing and healthcare[23], we developed an innovative analysis framework called *Profile Monitoring for Microbial Relationship Alteration* (PM2RA). This framework is designed to detect and quantify the relationship alteration within microbial community under different conditions. To our knowledge, PM2RA is the first method to make direct comparison of microbial associations between conditions, without initially constructing the co-occurrence network. By testing both synthetic and real datasets, we demonstrate that PM2RA is high in sensitivity and specificity, and that it identifies both previous and novel microbes in multiple kinds of disease. Moreover, PM2RA is robust in identifying important microbes in datasets obtained from different cohorts and different sequencing strategies. A web-based implementation of PM2RA is available at http://www.pm2ra-xingyinliulab.cn/.

## Method

### PM2RA methodology

We illustrated the PM2RA implementation framework in Figure. 1. The detailed framework is composed of following steps:

**Figure 1.**
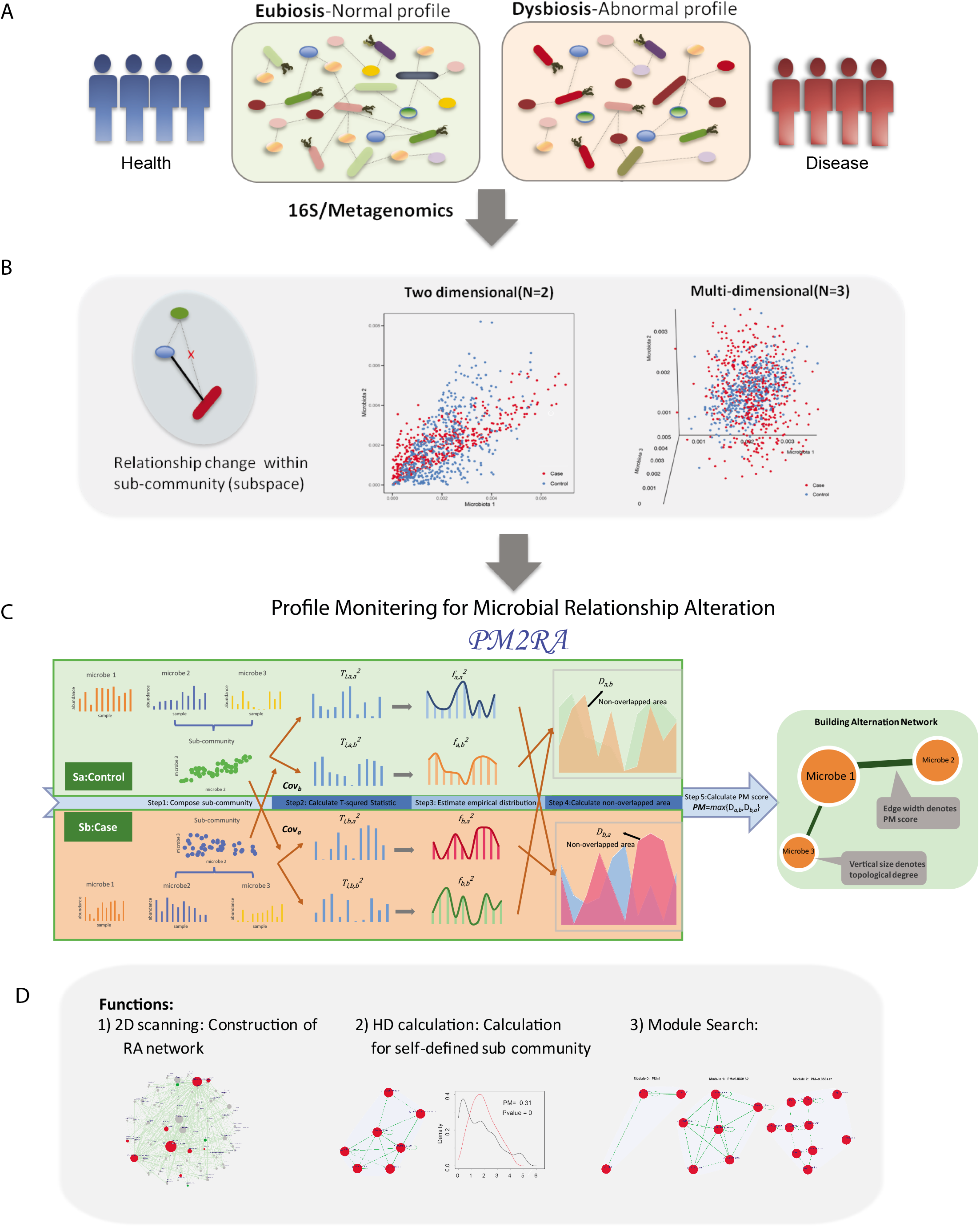
PM2RA methods. **(A)** In human disease, dysbiosis involves disturbed microbe relationships. (**B**) A relationship change can be involved in two or more microbes in a two-dimensional(2D) or highdimensional (HD) level. (**C**) PM2RA methodology framework. (**D**) The PM2RA was developed with three methods: 1) 2D scanning for pairwise association changes among the microbial community between two conditions; 2) High-dimensional (HD) calculation by which the PM score of any defined sub-community with two or more microbes could be calculated; and 3) a module search based on the HD calculation.

#### Step 1. Compose sub-community

In a microbiota profile, each two microbes and the interaction between them is a sub-community. In typical microbiome studies, the detected taxa or species ranges from tens to hundreds, so the relationship between two randomly chosen microbes is quite large. PM2RA qualifies all possible sub-community relationship alterations and output the alterations network.

#### Step 2. Calculate *T^2^* statistics

The Hotelling’s T^2^ statistics is one of the most popular statistics for monitoring the variables of a multivariate process[24]. This statistic takes into account both mean value and covariance matrix, which makes it suitable for reducing two or high dimensional microbial data into a one-dimensional data containing both abundance and relationship information. The *T*^2^ statistic is the multivariate counterpart of the t statistic and widely used in multivariate process consistence monitoring in both industry and biology fields[25,26]. It can be viewed as the generalized distance between the observed vector from the mean vector μ weighted by the inversion of covariance matrix, 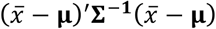 [27]. Since both **μ** and **Σ** are involved during the calculation, *T*^2^ statistic is sensitive to both relative abundance change and relationship change.

Let *S* = {*S*_a_, *S*_b_} denote the condition set between which microbe community changes are interested, e.g. *S* = {*S*_a_ = healthy, *S*_b_ = disease}. To guarantee symmetry, i.e. the community changes observed from condition *S_a_* to *S_b_* is equal to that from condition *S_b_* to *S_a_*, four types of *T^2^* statistic for each sub-community are calculated as following:

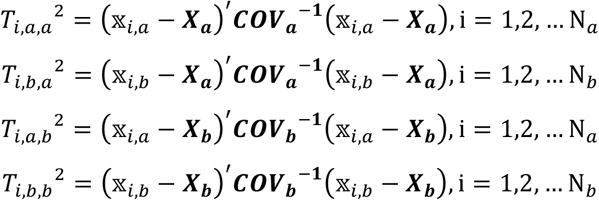

***X_a_, X_b_*** are the mean relative abundance vectors and ***COV_a_, COV_b_*** are covariance matrices of microbes under different conditions *S_a_* and *S_b_*. 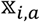 denote the microbe relative abundance of the i^th^ sample under *S_a_*. 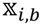 denote the microbe relative abundance of the i^th^ sample under *S_b_*.

In the pairwise relationship alteration analysis, the number of microbes is two and the sample size is usually much larger than that. In this case, and the possibility of strict collinearity between two microbes is very low, thus we can assume that the covariance matrix is nonsingular.

#### Step 3. Estimate empirical distribution of *T*^2^ statistics

Probability density function is an informative descriptive tool and can reflect the mean, standard deviation and other statistical properties of the dataset. To calculate alternations between two datasets, a straightforward way is to compare probability density functions of these two datasets. *T^2^* statistic follows a scaled chi-squared distribution under the assumption that samples have a normal distribution. This assumption is usually being violated in microbiota abundance context. Thus, PM2RA using kernel distribution to represent the probability density of *T^2^* statistics derived for each sub-community. Advantages of using kernel distribution is that the estimated kernel distribution produces a nonparametric, smooth, continuous probability curve that adapts itself to the data, rather than selecting a density with a particular parametric form (e.g. chi-squared distribution) and estimating the parameters. In this step, the kernel estimation method proposed by Scott, D. W[28] is applied on 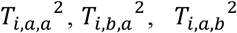 and 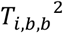. The estimated empirical probability density function is denoted as 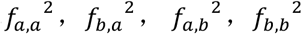 respectively. Outliers were removed from 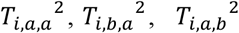 and 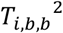 to get a robust estimation.

#### Step 4. Calculate the non-overlapped area between distributions

The non-overlapped area of two probability distribution functions is used to describe the difference between two sets of *T^2^* statistics.

The non-overlapped area of 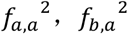 is

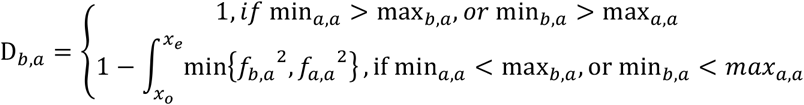

Where

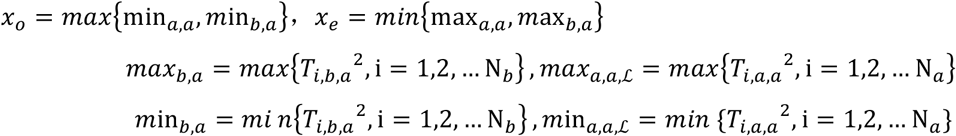

The non-overlapped area of 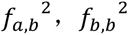 is

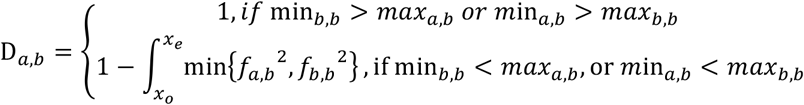

Where:

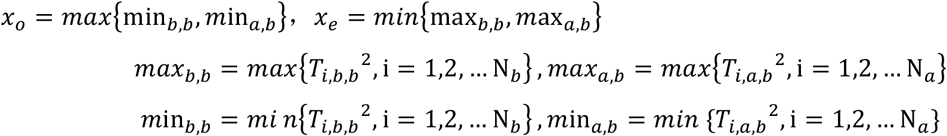

#### Step 5. Calculate PM score

The PM score is defined as = *max*{*D_ab_, D_b,a_*}. Compared with other non-parametric distance measures, e.g. Kullback-Leibler divergence, PM score has several advantages. The profile change measure is designed under symmetry. PM score has finite domains, i.e. [0,1], since the integration of a single probability density function is 1. The finite domains make PM score more interpretable. A Kolmogorov-Smirnov test is used on *T^2^* statistic to determine whether there is statistically significant difference between conditions.

#### Step 6. Construction of relationship alteration network

After traversing of all sub-communities, a weighted network is built to visualize overall relationship alternations (RA). In the RA network *G*=(*V,E*) where *V* is the set of vertices representing microbes and *E* is the set of edges denoting the relationship alteration between the two conditions. The edge width and vertices size denote PM score and topological degree, respectively.

### Identify the microbiome module with the largest alternation

In practice, the sub-communities with largest alterations between conditions are needed to guild microbial intervention for many diseases. These sub-communities may consist of two to more microbes as the interaction between microbes is not necessarily dual. In a microbial community, the total number of rational sub-community 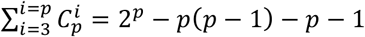 is an extremely large number when the number of microbes *p* grows. A greedy algorithm is designed to search subcommunity with large PM score as following.

1. Calculate profile changes for all sub-communities composed of two microbes.
2. Select N sub-communities with top PM scores and no overlapping microbes with each other. Mark the selected sub-communities as seed-communities.
3. Create a new sub-community set to be searched.

3.1 Adding a new microbe to a seed-community. This operation will generate new subcommunities whose dimension is dimension of the seed-community plus 1.
3.2 Combining two different seed-community. This operation will generate new sub-communities whose dimension is 2 times dimension of the seed-community.
4. Calculate PM scores for all sub-communities generated in step 3.
5. List all calculated PM scores results in step 1 and step 4.
6. Select N sub-communities with top PM scores and no joint microbe with each other based on the data list in step 5. Mark the selected sub-communities as new seed-communities. Loop into step 3 and start iteration.
7. When iteration time exceeds the pre-set threshold or the result is converged between two iterations, stop the searching process.
8. This algorithm will output N microbe modules.

### Dataset

Different types of datasets were downloaded to evaluate the performance of PM2RA. The 16s rRNA sequencing for CRC, overweight and obesity samples were downloaded from MicrobiomeHD[29]. The metagenomics data for CRC were download from[5]. The original dataset containing four cohorts from China, Australia, American, and Germany and French and America, while the America dataset, where the samples were collected more than twenty years ago, had no significant CA being detected (Table S1), thus we excluded it from our analysis. The diabetes datasets were downloaded from[30].

### PM2RA application

For each dataset, the relative abundance of microbes was calculated and those detected in less than 10% samples were removed from the following analysis. FDR<0.05 was used as a cutoff to filter significant RAs. The average computation time of PM2RA for a dataset contains 100 features 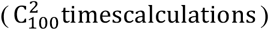 is 30 minutes (R with Parallel Computing on CentOS Linux release 7.6.1810 with E5-2680 v4, 8 cores).

### Synthetic microbial abundance datasets

In order to test the performance of PM2RA in realistic synthetic microbiome data, artificial datasets were generated based on the COMBO dataset[31], which contains OTUs from 100 samples. To evaluate the FPR of PM2RA, samples were randomly separated into two groups with an even sample size of 50, where it assumed that the relationship between any two OTUs does not change between the two groups. This process was repeated for 100 times to generate 100 matched artificial case and control datasets. PM2RA and other methods were applied to these datasets, and the FPR was calculated and compared. To evaluate the FNR of PM2RA, the abundance of any of the two OTUs were interchanged each time (for example A and B) to generate a case dataset where the abundance profile of the two selected OTU was changed, while others remained intact. In such a synthetic dataset, it assumed that the relationship between A/B and other intact OTUs will change in the case dataset, which is defined as the “True positive”. We interchanged the abundance of two microbes, which were not detected to be correlated by both SparCC and SPIEC-EAS, to generate the “case” datasets. The “control” dataset is the intact one. PM2RA and other methods were applied to these datasets, and the FNR were calculated and compared.

### Random Forest model

Random forest model (R 3.5.1, randomForest 4.6 package) were constructed based on the filtered microbe abundance (microbes detected in more than 10% samples) and T-square statistics of microbe pairs represented in the RA networks. The receiver operating characteristic (ROC) curve was obtained (pROC package) to evaluate the constructed models. One-sided *P-value* of AUC was assigned by bootstrapping (n=2000).

### Comparison with other methods

Two-step strategies were applied to all the synthetic datasets. First, the co-occurrence network for each synthetic case and control datasets were constructed using SPIEC-EASI and SparCC method implemented in the R package SpiecEasi (https://github.com/zdk123/SpiecEasi), the co-occurrence network was represented by a matrix consist of 1 (correlated) and 0 (not correlated). Second, the case and control co-occurrence network of each simulation were compared to obtain the changed association pairs. Lastly, the sensitivity and specificity of these methods were compared with that of PM2RA.

### Data availability

Source code of PM2RA and additional codes used in these work is available at https://github.com/bioinfolz/PM2RA.

## Results

### PM2RA framework

The human microbiome is a complex bacterial community where the relationships between microbes play important roles in the maintenance of eubiosis (Figure 1A). Examining the relationship alteration (RA) in microbiome between different healthy conditions provides additional insights into the pathogenesis of human disease (Figure 1B). PM2RA is specifically designed to quantify the RA involving two or multiple microbes (termed as sub-community) under different conditions. The basic idea of PM2RA analysis is to project the abundance data of two or multiple microbes under two conditions into the same space via Hoteling’s T2 statistics, and compare the difference in the distribution of T2 statistics to represent the RA between two conditions. A new scoring scheme called “PM score” is developed to quantify RA of each sub-community under different conditions through five steps which were described in the section of PM2RA methodology (Figure 1C). The more the relationship altered, the bigger PM score is. PM2RA quantify PM scores of all sub-communities. Next, a community alteration network was built in which edges denote the corresponding PM score (Figure 1C). Furthermore, the PM2RA has been developed with three functions (Figure 1D): 1) 2D scanning: scanning for pairwise microbe RA in the microbial community between two conditions. Based on the pairwise quantitative measurements, a RA network, can be constructed. Hub microbes in the RA network are those with extensively altered associations between two compared conditions; 2) High-dimensional (HD) calculation: in which the PM score of any defined sub-community referring to two or more microbes can be calculated; 3) Module search: based on the HD calculation, PM2RA can be applied to identify non-redundant sub-communities with maximum PM scores (termed as modules).

### Comparison on framework and Performance evaluation on synthetic data with other methods

In a traditional work flow (Figure 2A, left panel), the microbial co-occurrence network is constructed from the pairwise correlation, inverse covariance or other statistics based on microbial abundance in case and control samples respectively. The networks are then further compared by alignment-based or alignment-free methods. There are two drawbacks inherent in this pipeline. First, it is based on the pairwise correlation network constructed, but it is unclear whether correlation is a proper measure of association. Second, the association of microbiota is not necessarily dual; for example, multiple bacteria could form a tight community with weaker associations between any two members within it. Thus, the pairwise relationship analysis might ignore some functional associations consisting of multiple microbes. Furthermore, this comparison cannot quantify the association changes between conditions nor quantify the degree of difference between association changes. But rather, as shown in Figure 2A (right panel), PM2RA performs a direct comparison of relationship alteration among two or multiple microbes between conditions, no need to build co-occurrence network like that of the traditional method, and quantifies the RA as PM score.

**Figure 2.**
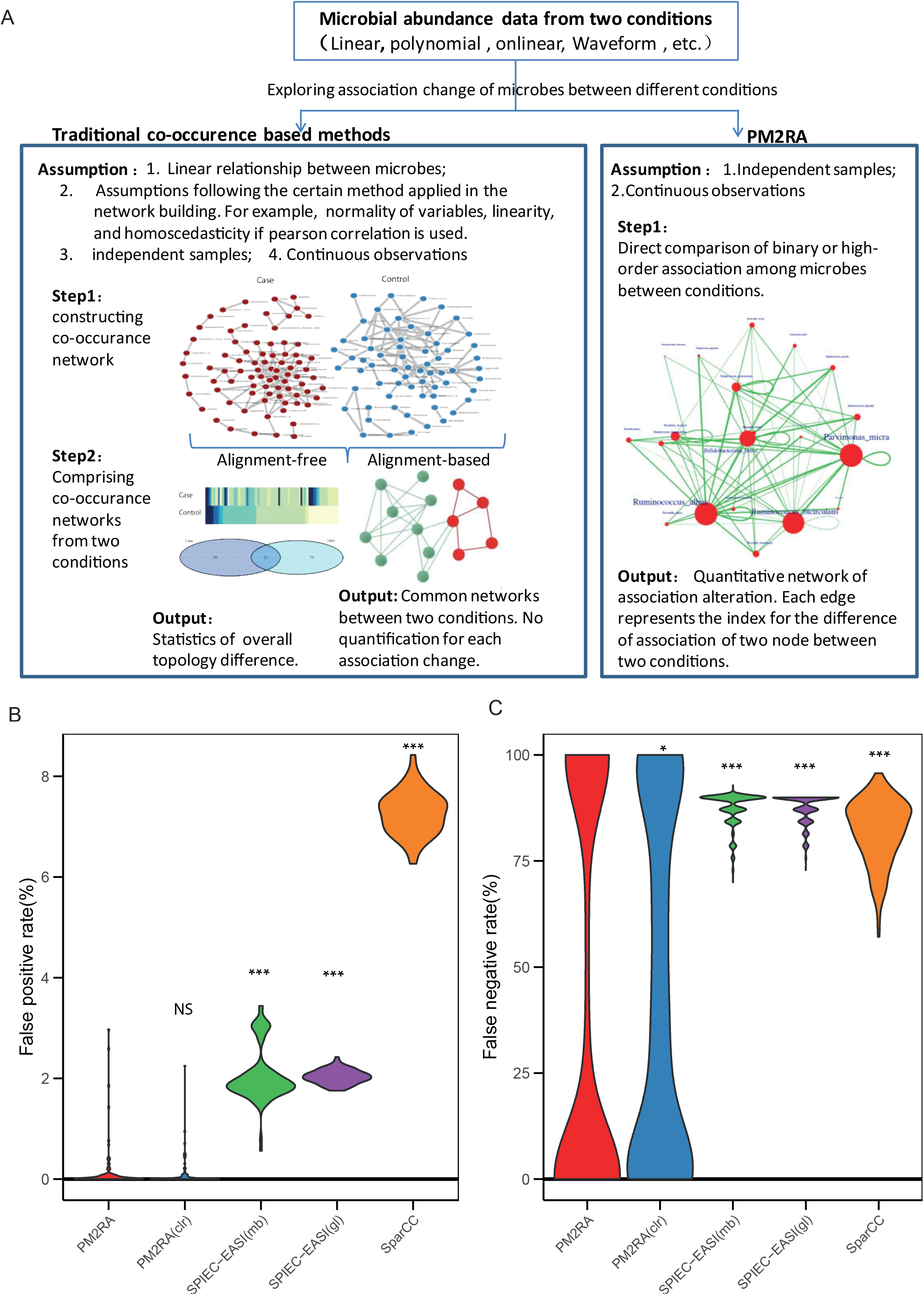
Comparisons between PM2RA and the co-occurrence-based methods. (**A**) The difference and advantages of PM2RA comparing to traditional co-occurrence-based methodologies. (**B**) FPR of different methods in detecting relationship changes. The methods illustrated from left to right are PM2RA applied to compositional data, centered-log-ratio transformed data, SPIEC-EASI (with either neighborhood selection method mb or covariance selection method glasso) and SparCC followed by a difference comparison. (**C**) FNR of different methods in detecting relationship changes. The difference was compared between PM2RA and other methods using the Mann-Whitney U test; *, **, *** correspond to P<0.05, 0.01 and 0.001, respectively.

To evaluate the performance of PM2RA in realistic synthetic microbiome data, the artificial datasets which contain operational taxonomic unit (OTUs) from 100 samples were generated based on a real microbiome dataset from the COMBO study[31]. First, the samples were randomly separated into two groups with an even sample size of 50, and this process was repeated 100 times to generate 100 matched artificial cases and control datasets. PM2RA and other two methods which were widely used to infer co-occurrence networks, i.e. SPIEC-EASI[32] and SparCC[33], were applied to these datasets. The average false positive rate (FPR, type I error) of PM2RA was significantly lower than the cooccurrence-based methods (PM2RA: 0.1%; mb-based: 2.1%; glasso-based: 2.0%; SparCC: 7.3%) (Figure 2B, Table S2), indicating a high specificity of PM2RA. Second, to evaluate the sensitivity of PM2RA, the synthetic datasets were generated by exchanging the abundance of two uncorrelated OTUs (Methods and Materials). PM2RA showed a significantly lower false negative rate (FNR, type II error) compared with the co-occurrence-based strategies (PM2RA: 33.5%; mb-based: 87.6%; glasso-based: 87.3%; SparCC: 82.1%) (Figure 2C, Table S3). The FNR of PM2RA is dumbbell-shaped (Figure 2C), suggesting that the FNR of PM2RA is affected by the effectiveness of the “case” datasets, and is sensitive to the correlations which were missed by SPIEC-EAS and SparCC (see Discussion section for details).

Compositional data analysis approach is widely used in microbiome data and has been proposed producing superior results in correlation analysis[34]. Therefore, to test the effect of compositional data on PM2RA performance, centered log-ratio (clr) transformation[34] was applied to the above artificial dataset. A similar FPR *(p-value=0.11)* was observed when applying PM2RA to the compositional and clr transformed data (Figure 2B). However, the FNR of applying PM2RA on compositional data was significantly lower than that of on clr transformed data (33.5% VS 43.4%) (Figure 2C). Taken together, the analysis showed that the compositional data was preferred in PM2RA to the transformed data.

### Robustness of PM2RA in colorectal carcinoma cohorts

To evaluate the performance of PM2RA in real microbiome data, a 16S rRNA sequencing dataset was obtained of colorectal carcinoma(CRC) [35], which is one of the key examples of complex diseases associated with dysbiosis of gut microbiota. A RA network containing 97 microbes and 607 significantly altered associations was identified (Figure 3A), and 12 genera with abundance changes were found to be involved in the RA network. The hub genera with the five largest degrees of topology were identified as *Porphyromonas*, *Parvimonas*, *Peptostreptococcus*, *Anaerostipes*, and *Dialister* (Figure 3A). The oral pathogens, *Peptostreptococcus*, *Porphyromonas,* and *Parvimonas*, were overrepresented in CRC, and accumulating evidence have shown that they promote the progression of oral cancer and other cancers of the upper digestive tract [36,37]. The other two hub genera, *Anaerostipes* and *Dialister* showed no difference in average abundance between the control and CRC groups (Figure 3B), but their associations with many other microbes were significantly altered (Figure 3A and Figure S1). *Anaerostipes spp.* is a butyrate-producing bacteria species that have been shown to play a key role in the maintenance of gut barrier functions. *Dialister* is reported to be overrepresented in oral cancer[38]. So, the analysis implicated that PM2RA can assistant us to find bacteria that affect CRC progression more accurately through searching for the CA between microbes.

**Figure 3.**
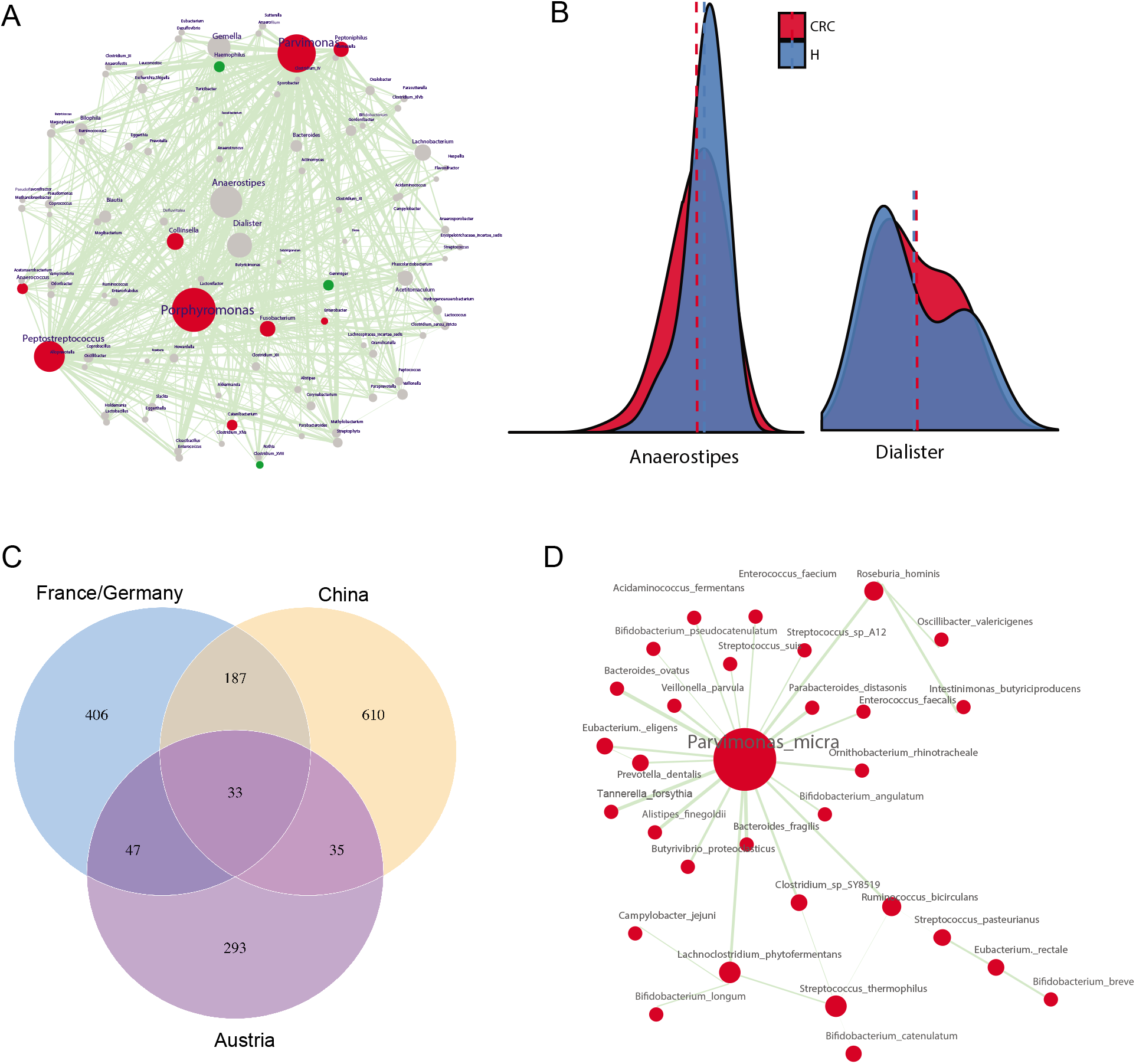
PM2RA detected common community alterations in different CRC cohorts. **(A)** The RA network for CRC. The node color represents the degree of difference between the case and control samples; red for microbes overrepresented in the CRC samples, green for microbes overrepresented in the control and gray for microbes not differentially represented. The size of nodes is proportional to their degree in the network, and the width of edges is proportional to the value of PM score. **(B)** No abundance difference between CRC and normal for *Anaerostipes* and *Dialister.* (**C**) Thirty-three association alterations were observed in all of the three CRC cohorts. (**D**) The common RA network among the three CRC cohorts.

The gut microbiome is highly dynamic and can be influenced by cohort-specific noise, and the results from differential abundance analysis may not be reproducible across different populations[39]. To investigate the robustness of PM2RA, metagenomics sequencing data of CRC patients and control subjects from Austrian, Chinese, and German/French cohorts[5] was tested (Figure S2A-C). Thirty-three common association changes were observed across the three cohorts (Figure 3C). Consistent with the results obtained from 16S rRNA sequencing data, *Parvimonas* spp. was identified as the top hub in the common RA network (Figure 3D). For example, the associations involving *Parvimonas micra* were extensively altered in the CRC group compared with the normal controls across the population. However, when measured by differential abundance, only three bacteria species were commonly detected in all three cohorts (Figure S2d), indicating PM2RA methodology is robust in identifying RAs.

### Robustness of PM2RA in metabolism disorders

We further assessed the robustness of PM2RA by investigating whether a common RA network can be observed in related diseases. PM2RA was applied to three closely linked metabolic disorders: overweight, obesity and diabetes. No significant association alterations or differential microbes were identified in the overweight cohort (normal: BMI<25; overweight: 25<=BMI<30), indicating BMI is not an informative index to assess a person’s disease state as has been previously reported[40]. In the obesity dataset (BMI >30), a RA network involving 85 altered associations and 97 bacteria genera were observed, with *Roseburia* having the most extensively altered associations with other genera (Figure 4A). Moreover, in diabetes cohorts A and B[41], there were 49 and 45 association changes involving *Roseburia spp.*, respectively. In diabetes cohort A, *Roseburia intestinalis* dominated the RA network (Figure 4B), while in diabetes cohort B *Bifidobacterium longum* was the top hub genus (Figure 4C). The clinical information showed comparable BMIs, but indicated a lower severity of dyslipidemia in diabetes cohort B. Studies have reported that *Bifidobacterium* spp. have anti-obesogenic or anti-diabetic potential[42]. The activated association changes with *Bifidobacterium longum* in diabetes cohort B may therefore be one explanation for the observed difference in dyslipidemia. By combining the three datasets, common association changes between *Roseburia* spp. and *Ruminococcus* spp. were identified, as well as changes between *Roseburia* spp. and *Bilophila* spp. (Figure 4D). *Roseburia* is one of the major butyrate-producing bacteria, and the modification in *Roseburia* spp. may affect various metabolic pathways[43]. In agreement with PM2RA analysis, animal experiments have demonstrated that *Roseburia* spp. can regulate the host immune system and reduce intestinal inflammation, which is also a marker for obesity and metabolic dysfunctions[44,45].

**Figure 4.**
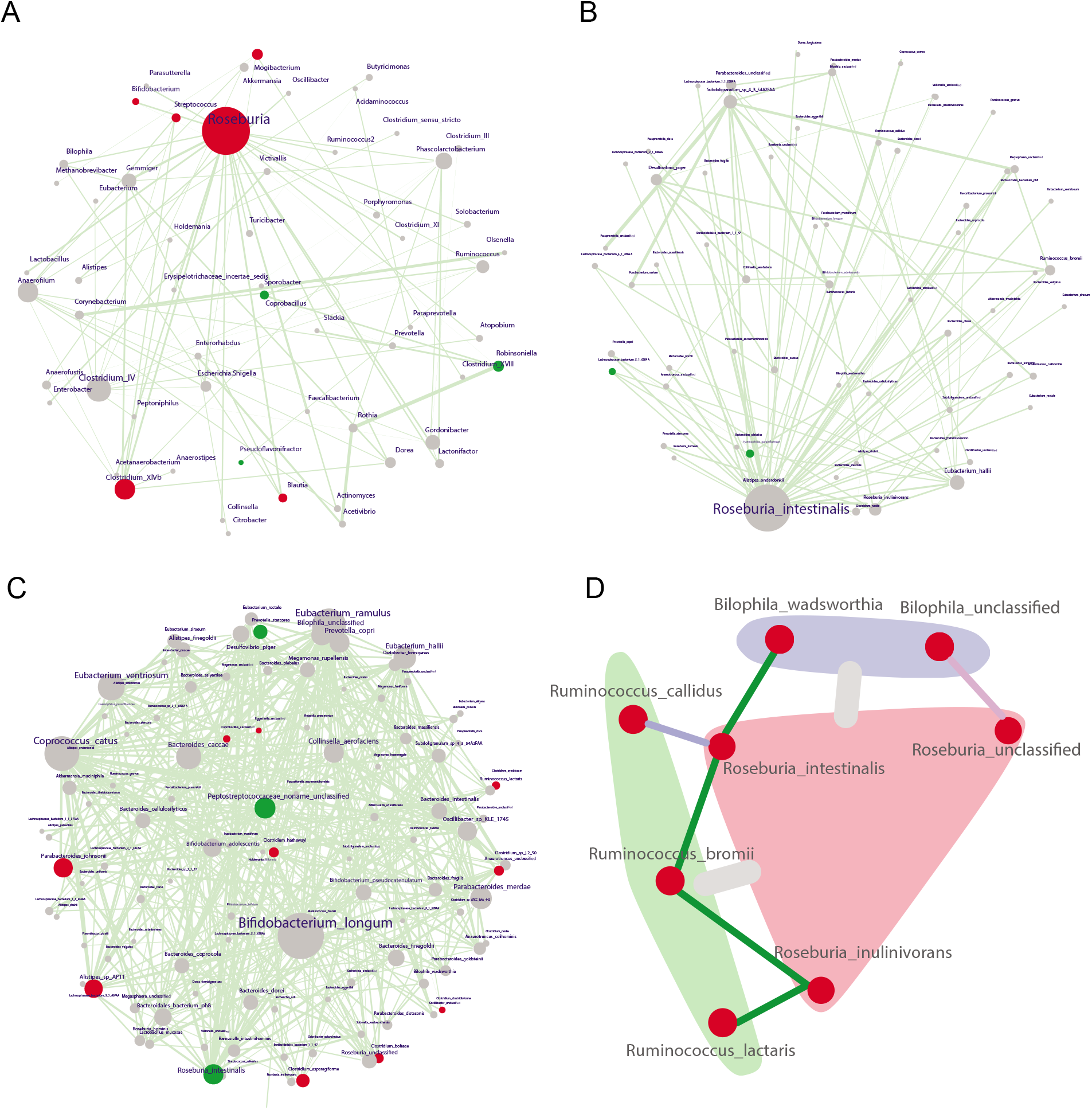
PM2RA detected common community alterations in multiple metabolic diseases. (**A**) The community alteration network for obesity (case=193, control=451). (**B-C**) The community alteration network for diabetes cohorts A (case=57, control=79) and B(case=99, control=99). **(D)** The common association alteration observed in obesity and diabetes datasets. At the genus level, the association between Roseburia (colored as pink module) and Ruminococus(colored as green module), and that between Roseburia and Bilophila(colored as purple module) were generally changed. The red, green and blue lines between species represent the association change were observed in diabetes cohort A, B and both, respectively.

Taken together, the consistent results obtained using different sequencing strategies and different cohorts indicate the robustness of PM2RA in identifying RA networks in various diseases.

### Comparing PM2RA with Netshift in real dataset

As above, PM2RA is an innovative method aiming to measure RA in microbial community directly. However, a few additional methods, for example, Netshift was designed to solve similar problems. Therefore, we further compared the results of PM2RA and Netshift in the real datasets.

Netshift is a co-occurrence-based method developed to quantify rewiring and community changes in microbial association networks between a state of health and one of disease[46]. It was designed to produce a score identifying important microbial taxa that serve as “drivers” from a state of healthy to one of disease. Netshift was applied to the dataset of colorectal carcinoma (Figure S3A-d) and metabolism disorders (Figure S4A-d) as mentioned previously. Two common driver species were identified in colorectal carcinoma (Figure S3E), i.e. *Butyrivibrio proteoclasticus* and *Streptococcus pyogenes.* However, the previously identified microbes involved in the disease, e.g. *Bacteroides fragilis, Fusobacterium nucleatum, Porphyromonas asaccharolytica, Parvimonas micra, Prevotella intermedia, Alistipes finegoldii, and Thermanaerovibrio acidaminovorans[47],* were not captured. However, five of the seven species were commonly detected by PM2RA in three colorectal carcinoma datasets (Figure 3D). More than 50% of the drivers identified by Netshift were shared by obesity and overweight samples (Figure S4E) and four drivers were shared by the two diabetes datasets (Figure S4F), i.e. *Alistipes shahii, Anaerotruncus colihominis, Eubacterium hallii,* and *Eubacterium ventriosum*. However, few of these drivers had been previously reported as to be associated with metabolism disorders. For example, the repeatedly reported species, *Ruminococcus. sp.* and *Roseburia. sp.,* were detected by PM2RA (Figure 4D), were not detected as drivers by Netshift.

### PM2RA is more effective in distinguishing case and control samples

To test whether the microbial relationship represented in PM2RA (the Hoteling’s *T^2^* statistics) captured important information that distinguishing cases from control samples, we generated random forest (RF) models using the total microbe abundance (RF-A), microbe abundance of drivers detected by Netshift (RF-N) and the Hoteling’s T^2^ statistics (RF-P) of each pair of microbes in the RA networks in the above datasets, respectively. The RF-P models achieved AUC values of the ROC curve higher than that of RF-A models in 6 of the 7 datasets (Figure 5). In the 16S rRNA CRC and Diabetes cohort, the AUC value of RF-P models was significantly higher than that of RF-A models (*p-value* =0.012 for CRC and *p-value* =0.017 for diabetes cohort A). In the comparison of RF-P and RF-N, significantly increased AUC values were observed in the entire cohort (Figure 5). These results indicated that the RA revealed more information than abundance shift of the whole microbiome as well as the abundance shift of drivers identified by the co-occurrence-based method in the pathogenesis of these diseases. In addition, the hub microbes in the RA network were highly overlapped with the microbes with the highest importance scores in the RF-A model (Figure 3–4 and Figure S5-6). In the 16S rRNA CRC dataset, the top three hubs microbes, i.e. *Parvimonas, Porphyromonas* and *Peptostreptococcus* (Figure 3A), were ranked as the top four important features in the RF-A model. And the *Parvimonas* species, the most notable hub microbe commonly detected in multiple CRC datasets (Figure 3D), was among the top 5 most important species in three out the four CRC cohorts (Figure S5A-D). The *Roseburia* and *Bilophila* species which were commonly detected by the PM2RA in obesity and diabetes (Figure 4D) were identified by RF-A model as top important features (Figure S6A-C). However, the *Ruminococcus* species identified by PM2RA was not recognized as a top important feature in the RF-A model, which may represent the additional information captured by PM2RA that contributed to the higher classification power. These results suggested that the Hoteling’s *T^2^* statistics transformation in PM2RA not only preserved the most important feature that distinguishing health and disease status, but also provided extra information that underlying the pathogenesis of human diseases.

**Figure 5.**
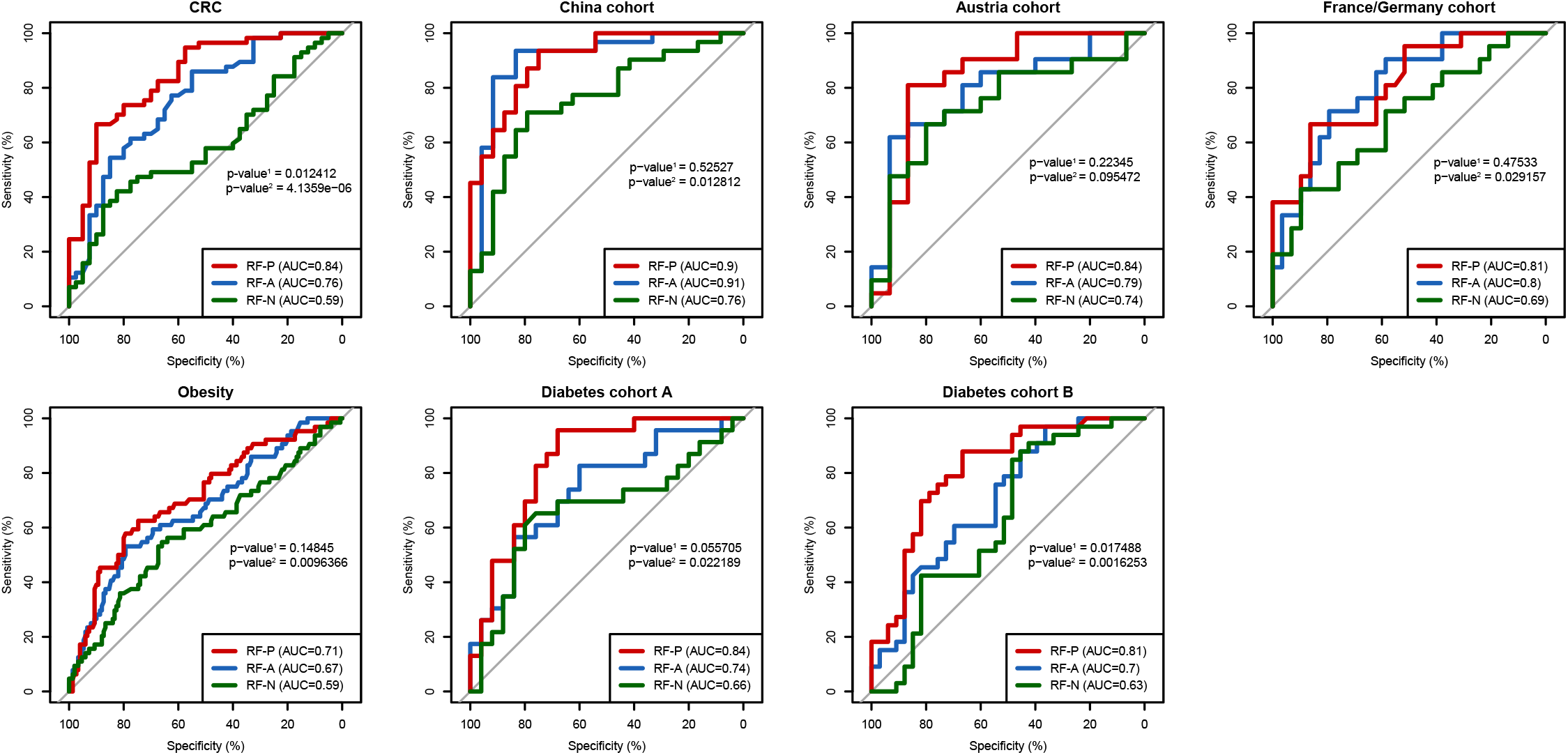
The ROC curve of the RF models revealed PM2RA is more predictive. The p-value^1^ and p-value^2^ denote the p-value for comparison between Hoteling’s *T^2^* statistics (RF-P) with total microbe abundance (RF-A) and RF-P with microbe abundance of drivers detected by Netshift (RF-N), respectively.

### High dimensional PM2RA analysis is complementary to 2D scanning

Associations between microbes are not necessarily structured in a paired way, and multiple microbes are able to form closely interacting sub-communities. The ability of PM2RA to quantify RA involving multiple microbes makes it applicable to identifying alterations in such communities. We therefore tested the performance of PM2RA in high-dimensional microbial communities in the abovementioned datasets. A greedy algorithm was designed to search for RA networks with the highest PM score in multiple dimensions. By applying the greedy algorithm, high dimensional RAs (HD-RA, FDR <0.05, PM score >0.6) were identified in all datasets except for the obesity and overweight datasets (Table S4). Most modules contained more than two microbes, indicating potential associations among multiple bacteria. Furthermore, many HD-RAs contained microbe pairs that were not significantly altered at the 2D-level (Figure 6A-B), illustrating the increased ability of PM2RA to detect weak change signals in HD microbe communities under different conditions, which have usually been ignored by 2D scanning (Figure 6C). These results suggested that PM2RA was a promising method to quantify two and high-dimensional microbial community alteration.

**Figure 6.**
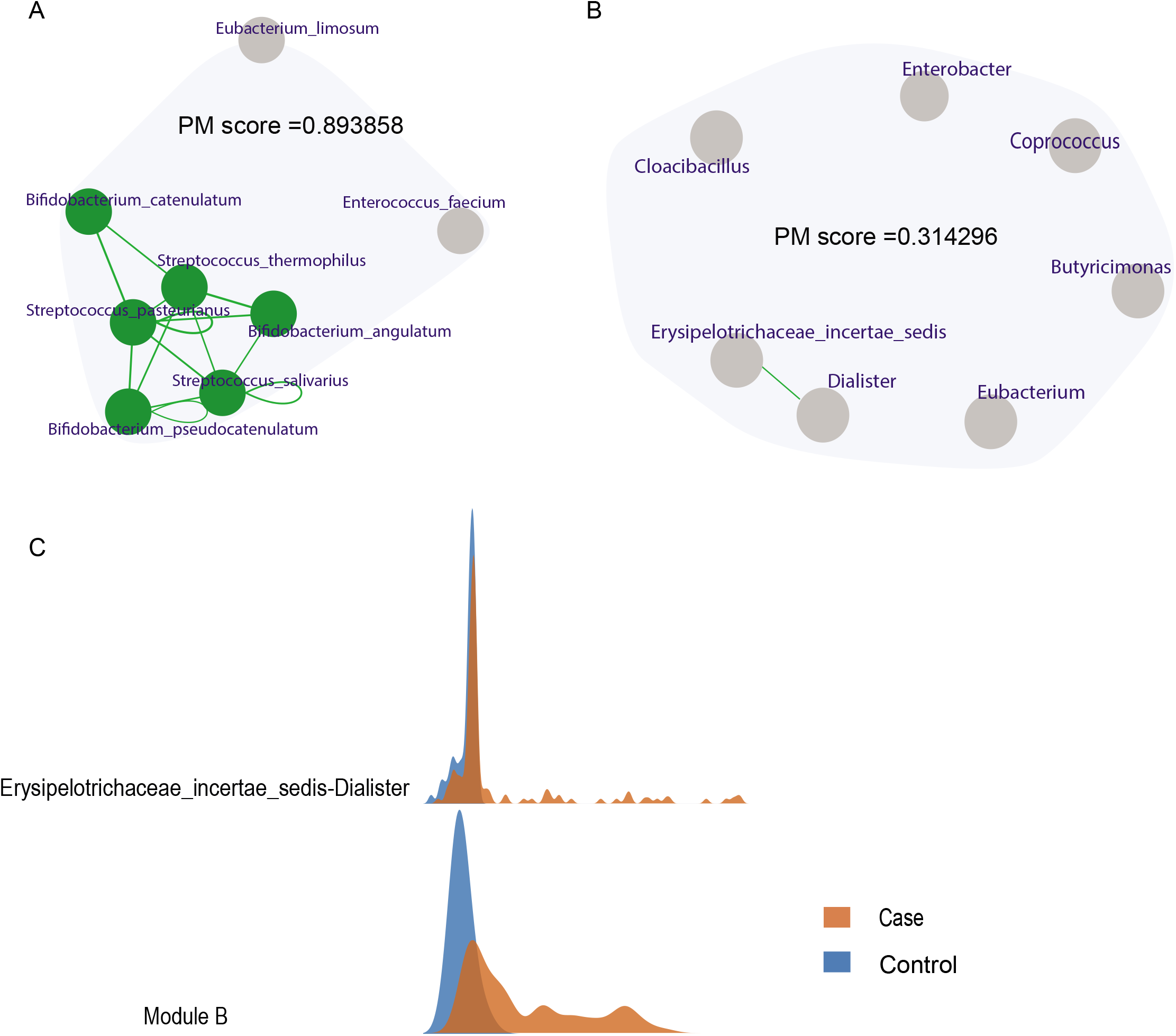
High dimensional PM2RA analysis is complementary to 2D scanning. **(A-B)** Examples of high dimensional RAs. The lines between microbes represent the RA detected with 2D scanning. (**C**) A comparison of PM scores for two-dimensional (Erysipelotrichaceae incertae sedis and Dialister) and high dimensional module in **B**.

## Discussion

Microbial association analysis is an important complement to abundance differential analysis in the study of dysbiosis in disease. In the current study, we developed an innovative analysis method to detect and quantify microbial relationship alternations. PM2RA measures the relationship alteration of microbial sub-communities, without initially constructing a correlation network for each condition. Compared to the traditional workflow from the occurrence-based network, we demonstrated that PM2RA is higher in sensitivity and specificity. And random forest analysis revealed that the relationship represented by PM2RA is more predictive for distinguishing disease and health status compared with the microbe abundance and abundance of driver microbe identified by co-occurrence-base method. Furthermore, application of PM2RA in several disease datasets demonstrated the robustness of PM2RA.

In our applications, PM2RA showed biological reproducible results in datasets with samples size ranging from tens to hundreds. However, since PM2RA calculates difference based on the projection distribution, the larger the sample size, the more precise the distribution estimation, and so we recommend applying it to dataset with more than 30 samples for each of the compared conditions.

When evaluating the sensitivity of PM2RA, defining the true positive relationship alteration was difficult, due to the lack of statistical methods to quantify and defining relationship change. We used SPIEC-EAS25 and SparCC26 to defined “uncorrelated” microbes, and interchanged the abundance of any two uncorrelated microbes to generate the “case” datasets. However, some types of correlation can still be neglected, thus rendering the exchange not fully effective. Therefore, the results might have shown an underestimated sensitivity. The false negative rate (FNR) of PM2RA is dumbbell-shaped (Figure 2C), which suggesting that the FNR of PM2RA affected by the effectiveness of the “case” datasets. When the exchanged OTUs are independent from each other, a very low FNR will be observed. Otherwise, PM2RA will recognize the relationship between most other species with them as similar, resulting in a high FNR. These results also indicated that PM2RA sensitive to the correlations which were missed by SparCC and SPIEC-EAS.

The abundances of microbial OTUs from amplicon-based datasets are usually compositional; counts are normalized to the total number of counts in the sample. Applying traditional correlation analysis to such data may produce spurious results[34]. Since PM2RA detects RA without constructing cooccurrence, the influence of compositional data on results is small. Therefore, a comparable specificity was observed when applying compositional data and centered-log-ratio (clr) transformed data to the synthetic datasets. However, the sensitivity of applying PM2RA on clr-transformed data was significantly lower than that of on compositional data. This might have been due to the transformation’s alteration of the abundance baseline and the subsequent impact on the relationship inherited in the raw abundance data.

In conclusion, PM2RA is a novel method for identifying and directly quantifying the relationship change in microbial communities. It circumvents the drawbacks of the co-occurrrence-based methods, and the applications to multiple human diseases reveal biologically significant results. The ability of PM2RA to detect community-level dysbiosis may become a useful tool for exploring the functional alterations of microbes as a whole in a variety of diseases or biological conditions, and further provide additional hints about the pathogenesis of human disease.

## Author Contributions

X.L and K.M. conceived this study. X.L, K.M, Z.L and Z.X. design this study. K.M. and Z.L. developed the methodology. Z.L constructed the online service and local packages. Z.L, K.M, X.L and Q.Z prepared figures and statistical analysis. Z.L., X.L, Z.X. and K.M. wrote the manuscript. All authors read and approved the final manuscript.

## Acknowledgments

This work was supported by NSFC grant 81671983 and 81871628, Natural science funding BK20161572 from Jiangsu province and starting package from NJMU (X.L.). Starting funding for the team of gut microbiota research in NJMU (X.L.)

## Supplementary Figures

**Supplementary Figure 1** (**A,B**) Association changes involved by Dialister (A) and Anaerostipes (B).

**Supplementary Figure 2** The RA network for CRC cohorts from China (**A**) (case=73, control=92), France/Germany (**B**)(case=88, control=64) and Austria (**C**) (case=46, control=63). The node color represents the degree of difference between the case and control samples; red for microbes overrepresented in the CRC samples, green for microbes overrepresented in the control and gray for microbes not differentially represented. The size of nodes is proportional to their degree in the network, and the width of edges is proportional to the value of PM score. **(D)** The overlap of differential represented microbes across three CRC cohorts.

**Supplementary Figure 3 (A-D)** The drives identified by “Netshift” method in colorectal carcinoma.**. (E)** The overlap of drives across three CRC cohorts.

**Supplementary Figure.4** The drives identified by “Netshift” method in metabolism disorders.(**A**) overweight; (**B**) Obesity;(**C**) Type 2 diabetes dataset A; (**D**) Type 2 diabetes dataset B; (**E**) The overlap of drives between over-weight and obesity. (**F**) The overlap of drives between two diabetes datasets.

**Supplementary Figure 5** The top 20 important features by RF model generated on the microbe abundance in four CRC cohorts. The red labeled microbes are taxa identified by PM2RA as hubs of the RA network.

**Supplementary Figure.6** The top 20 important features by RF model generated on the microbe abundance in the obesity and diabetes cohorts. The red labeled microbes are taxa identified by PM2RA as hubs of the RA network.

## Supplementary Tables

**Supplementary Table 1**. False positive rate of PM2RA and other methods in synthetic datasets.

**Supplementary Table 2**. False negative rate of PM2RA and other methods in synthetic datasets.

**Supplementary Table 3**. Microbe modules in each dataset.

**Supplementary Table 4**. 2DScanning results for America CRC.

