## Supplementary material for "A framework for detecting and quantifying relationship alterations in microbial community: Quantifying microbial relationship alteration": Figure S1; Figure S2; Figure S3; Figure S4; Figure S5; Figure S6

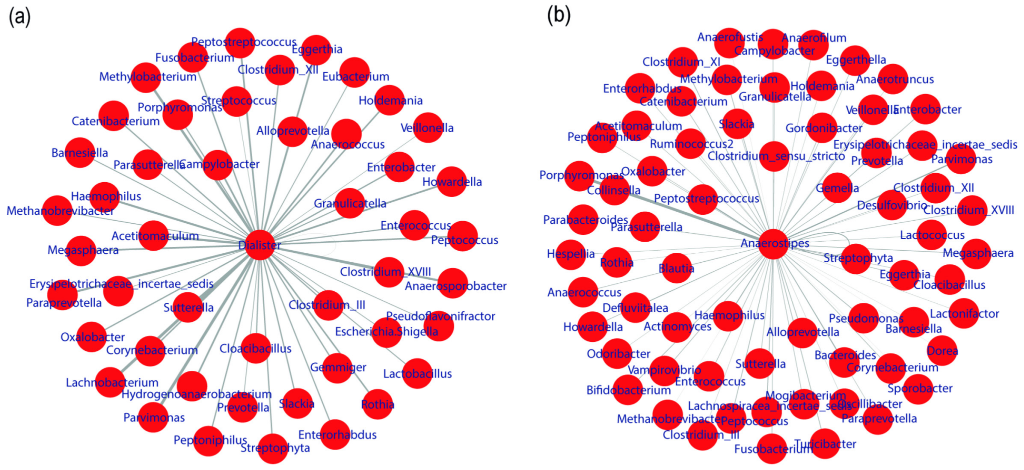


**Supplementary Figure 1** **(a)** Fourteen microbes differed significantly between CRC patients and controls. The *** and ** denote FDR < 0.001 and FDR < 0.05, respectively. **(b,c)** Association changes involved by Dialister (b) and Anaerostipes (c).

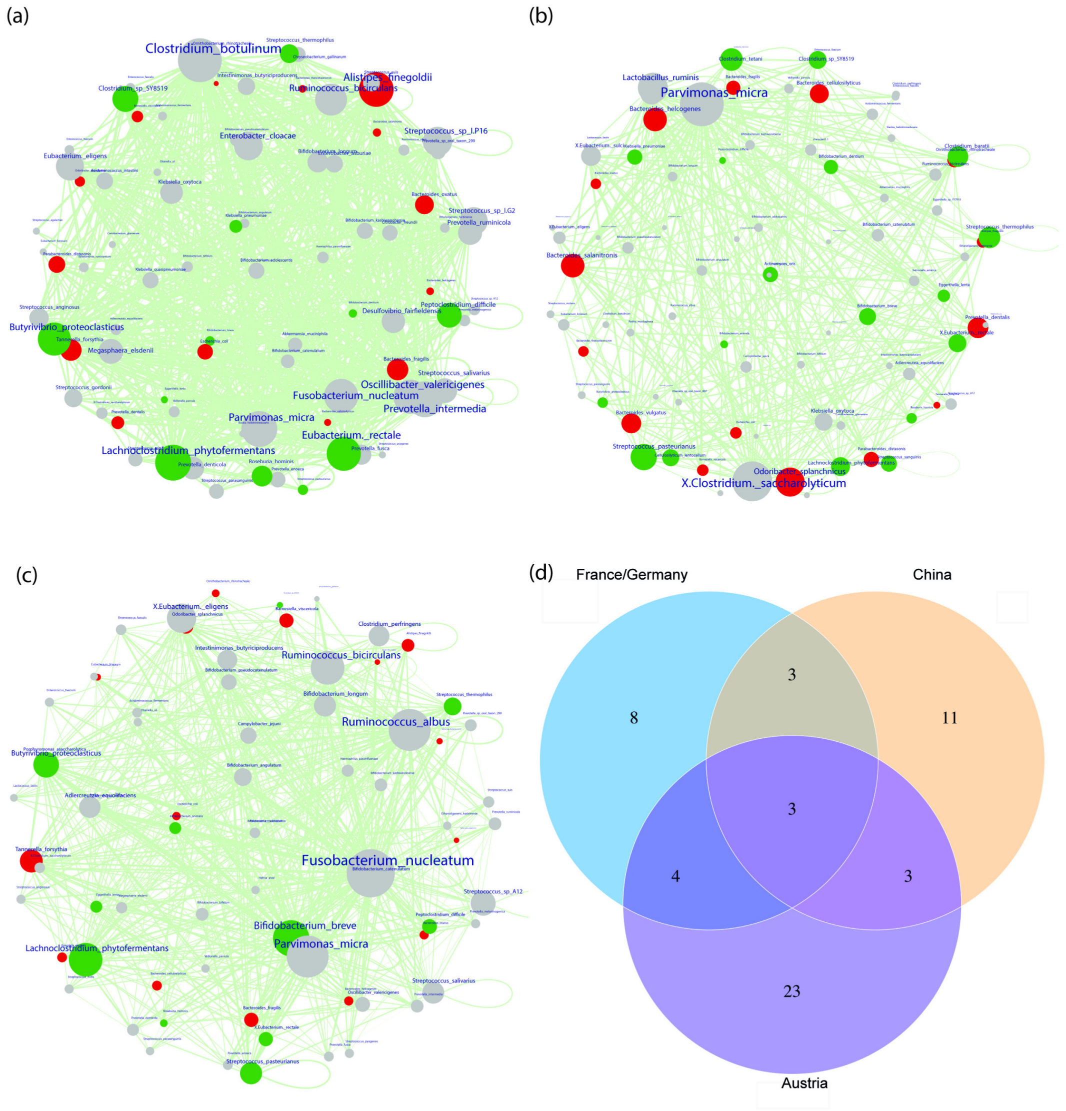


**Supplementary Figure 2** The RA network for CRC cohorts from China (**a**) (case=73, control=92), France/Germany (**b**) (case=88, control=64) and Austria (**c**) (case=46, control=63). The node color represents the degree of difference between the case and control samples; red for microbes overrepresented in the CRC samples, green for microbes overrepresented in the control and gray for microbes not differentially represented. The size of nodes is proportional to their degree in the network, and the width of edges is proportional to the value of PM score. **(d)** The overlap of differential represented microbes across three CRC cohorts.

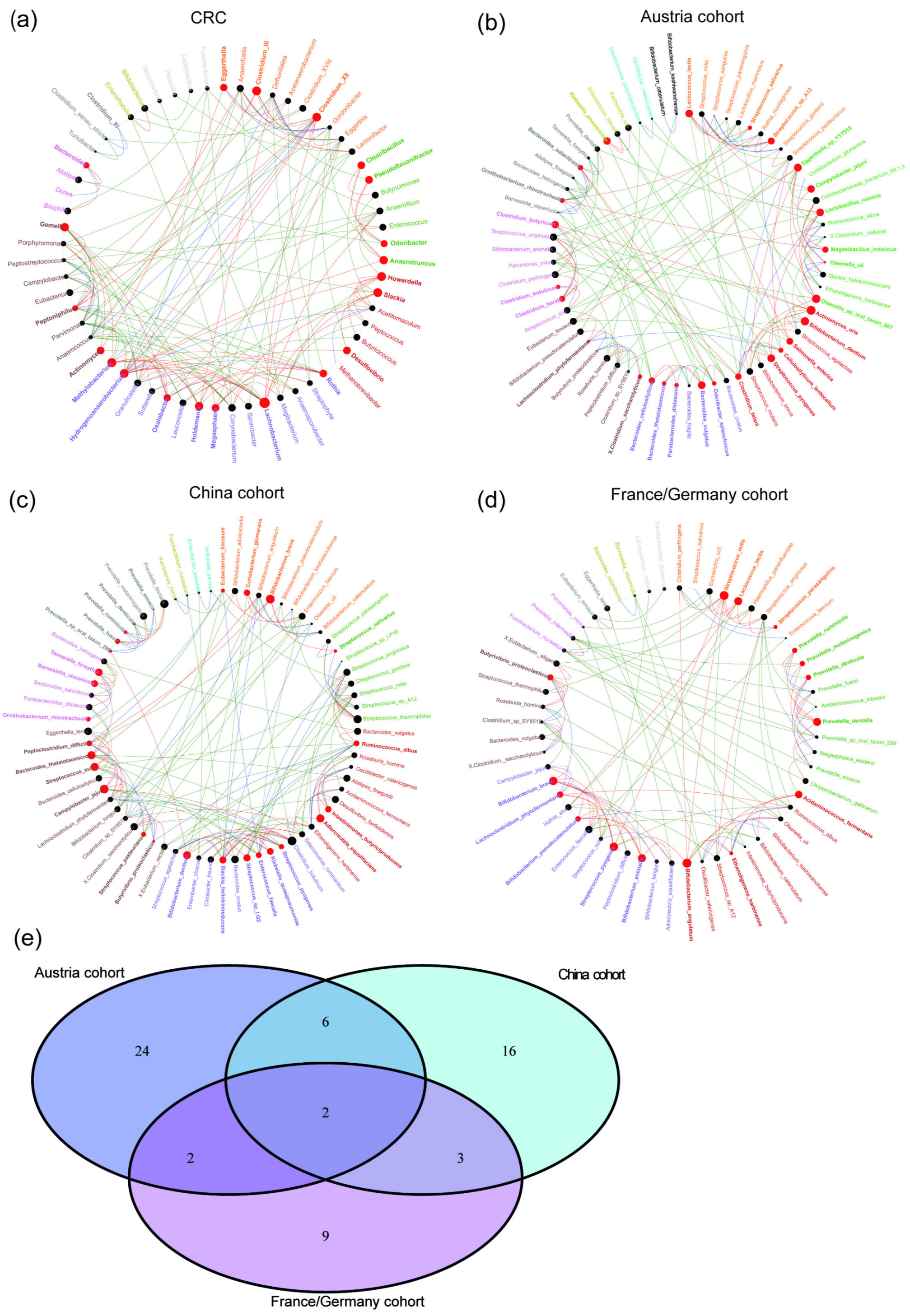


**Supplementary Figure 3 (a-d)** The drives identified by "Netshift" method in colorectal carcinoma**. (e)** The overlap of drives across three CRC cohorts.

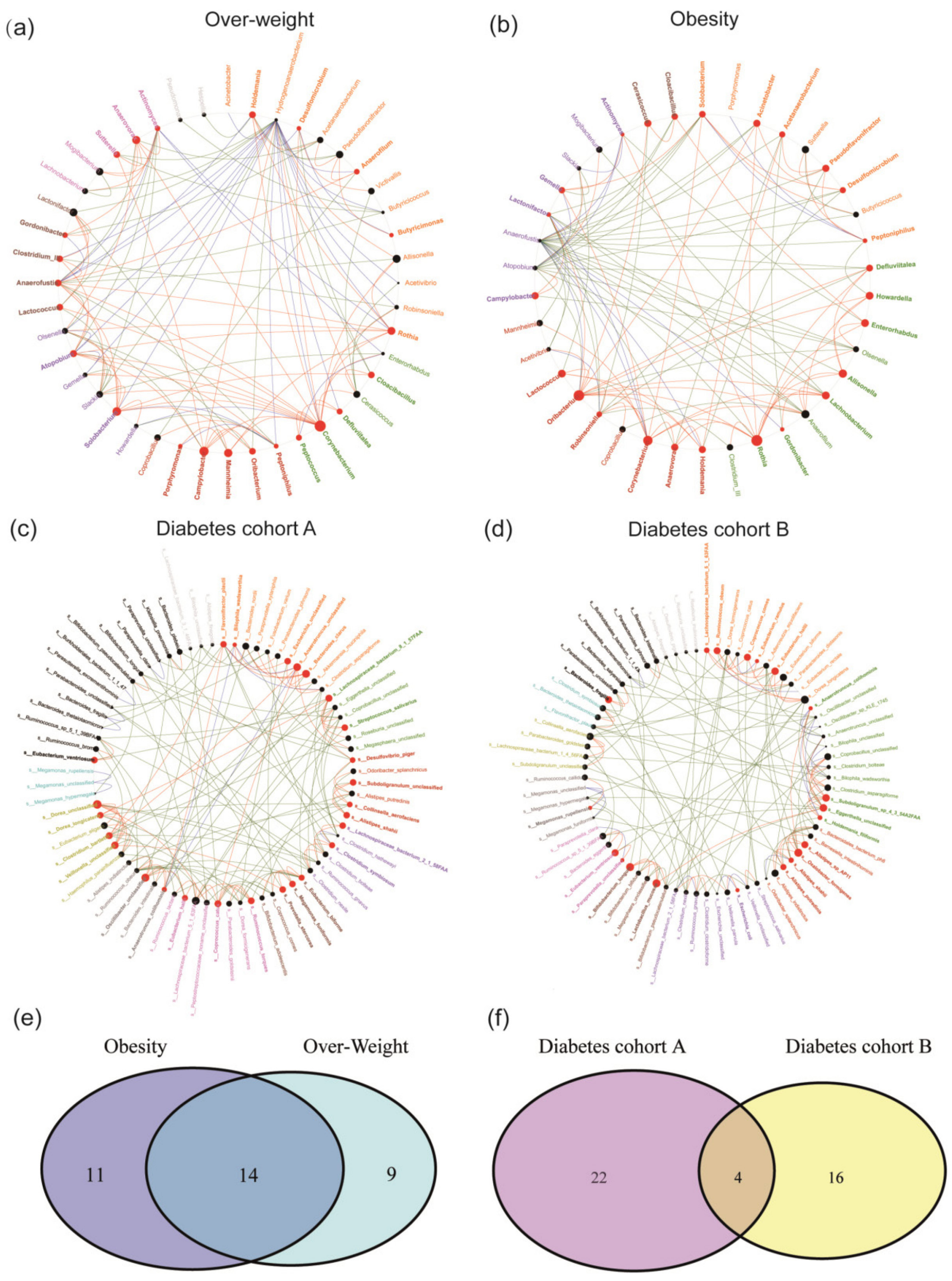


**Supplementary Figure 4** The drives identified by "Netshift" method in metabolism disorders. (**a**) overweight; (**b**) Obesity; (**c**) Type 2 diabetes dataset A; (**d**) Type 2 diabetes dataset B; (**e**) The overlap of drives between over-weight and obesity. (**f)** The overlap of drives between two diabetes datasets.

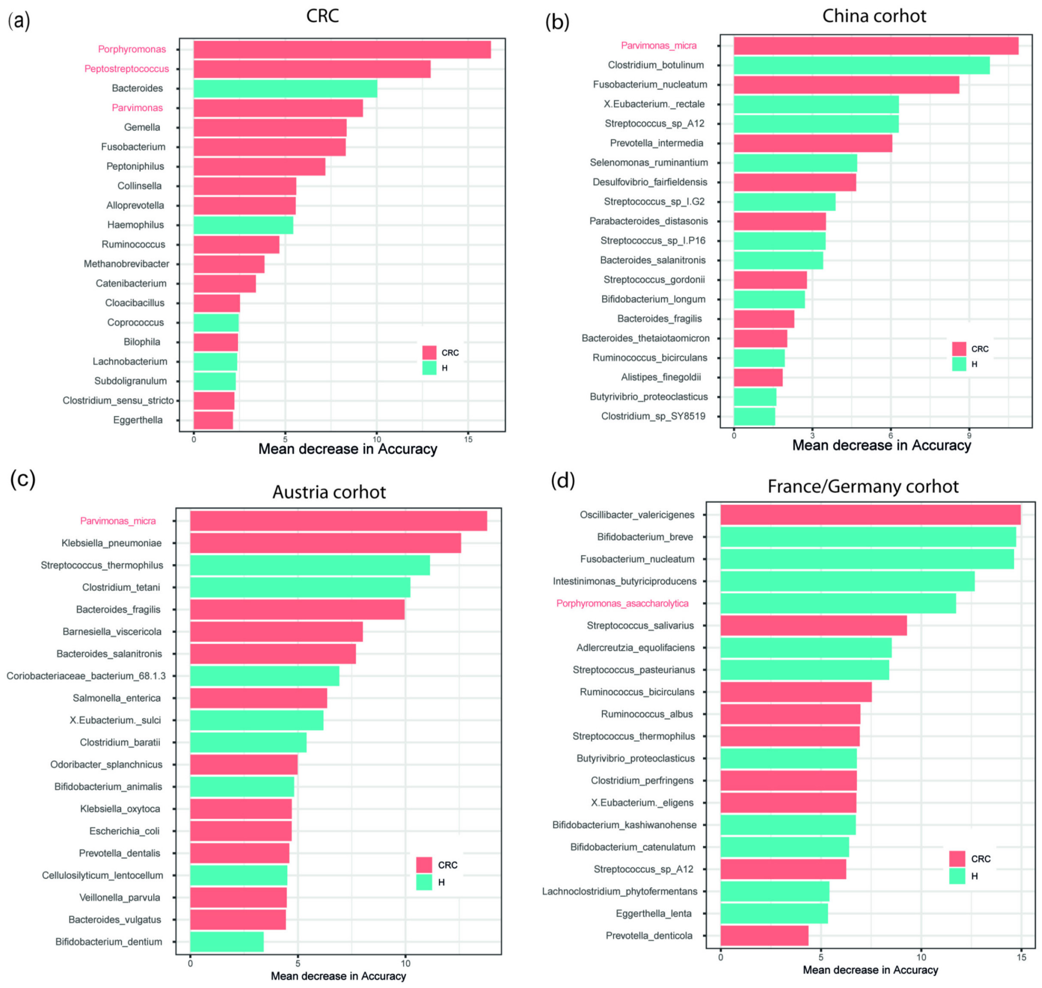


**Supplementary Figure 5** The top 20 important features by RF model generated on the microbe abundance in four CRC cohorts. The red labeled microbes are taxa identified by PM2RA as hubs of the RA network.

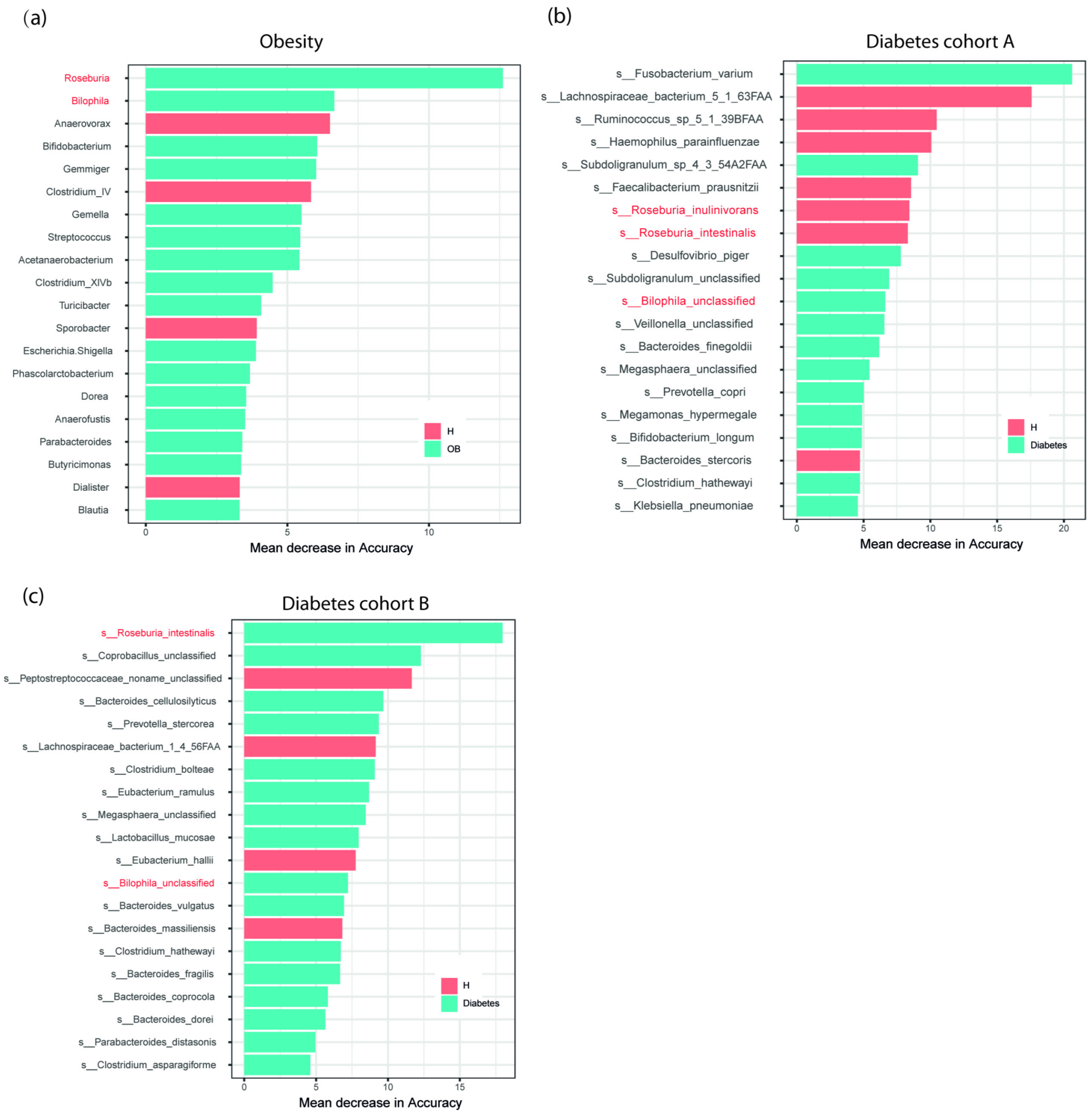


**Supplementary Figure 6** The top 20 important features by RF model generated on the microbe abundance in the obesity and diabetes cohorts. The red labeled microbes are taxa identified by PM2RA as hubs of the RA network.
